# The effect of delayed processing on ovarian tissue stored for fertility preservation

**DOI:** 10.1101/2020.06.30.180190

**Authors:** M S Zemyarska, B D Bjarkadottir, X Wei, C A Walker, S M Lane, J Davies, S A Williams

**Affiliations:** Nuffield Department of Women’s and Reproductive Health, University of Oxford, Women’s Centre, John Radcliffe Hospital. Oxford, United Kingdom. OX3 9DU; Department of Paediatric Oncology and Haematology, Children’s Hospital Oxford, Oxford University Hospitals NHS Foundation Trust, Oxford, United Kingdom. OX3 9DU; Oxford Cell & Tissue Biobank, Children’s Hospital Oxford, Oxford University Hospitals NHS Foundation Trust, Oxford, United Kingdom. OX3 9DU

**Keywords:** fertility preservation, delayed processing, ovarian cryopreservation, follicle, human, sheep, conservation

## Abstract

**Background:** Ovarian tissue cryopreservation (OTC) is important for fertility preservation and conservation. Delay in OTC may be required for transport or workflow management, however little is understood about the effect of processing delay on the tissue.

**Objective:** To determine whether a delay of 24-48 hours to OTC affects primordial follicle (PF) health.

**Methods:** Ovaries (n=6 sheep) were processed immediately or after storage at 4°C (24h, 48h). Tissue was fixed fresh, after cryopreservation or 10-day xenotransplantation. Morphological assessment of follicle health and development was performed.

**Findings:** A total of 1541 follicles were analysed. A 24h processing delay did not impact PF health in fresh or cryopreserved tissue. In fresh tissue a 48h delay had an adverse effect on follicle health (OR=2.47, 95% CI 1.29-4.71). Interestingly, a 48h delay resulted in cryopreserved tissue being less likely to be graded as unhealthy compared to control (OR=0.56, 95% CI 0.36-0.87). There was no difference in PF health or development across groups following xenotransplantation.

**Conclusion:** Ovarian tissue can be stored for up to 48 hours prior to cryopreservation with no net impact on PF health.

## Introduction

At present, over 80% of children and adolescents diagnosed with cancer will survive the disease and reach adulthood (1). Furthermore, as of 2011 there are an estimated 388,000 childhood cancer survivors in the US (2) and over 30,000 in the UK (3). Childhood cancer treatment can, however, often be associated with a significant negative impact on the growth, development and, commonly, the fertility of young patients (4). Thus, fertility preservation is of importance to young oncology patients, in order to prevent further emotional distress at the time of cancer diagnosis and long-term negative impact on the recovery and future wellbeing of childhood cancer survivors (5).

Ovarian tissue cryopreservation is a method of fertility preservation that is becoming increasingly well-established world-wide. This involves dissection, cryoprotection and freezing of the ovarian cortex with the aim of preserving the primordial pool of follicles contained within. Importantly, although the intervention is still considered experimental in several countries, as of 2017, an estimated 130 pregnancies have been achieved after ovarian tissue transplantation globally (6). However, ovarian tissue cryopreservation remains a specialised procedure, which requires practical experience, specialist equipment and dedicated facilities, which comply with EU and/or national regulations. As a result, although individual experimental cases have been reported from 21 countries (7), the availability of the technique as a modality of fertility preservation is limited (8), while the need for it may continue to rise.

One suitable approach to improve accessibility to ovarian tissue cryopreservation involves ovarian tissue being procured at various satellite clinics and transported to a central cryopreservation facility; this has been termed the ‘Danish model’ (8,9). However, the Danish model may require overnight storage of the tissue, to allow for long-distance transportation of the samples and co-ordination and workload management at the cryopreservation facility. This may create a delay between ovarian tissue retrieval and processing for cryopreservation, during which the ovary is in a state of ischaemia, which can cause nutrient deprivation, hypoxia and, as a result, cell death (10). The process of tissue cryopreservation and thawing is further associated with great physiological stress to living cells and tissues due to the dramatic changes in osmolarity and temperature experienced by the tissue, as well as the formation of structurally disruptive ice crystals (11). Hence, it is of concern that pre-exposure of the ovarian tissue to challenging conditions such as prolonged cold ischaemia has the potential to exacerbate the damaging effects of freeze-thaw on tissue health and integrity, even if no direct effects were observed in fresh tissue.

At present, two case reports of cryopreservation delay for up to 20 h at 4°C with one subsequent live birth have provided proof of principle that follicles can survive freeze-delay and that ovarian cortical tissue can retain graft functionality (8,12). However, the impact of cryopreservation delay on the health of the primordial pool remains controversial, with studies reporting either no effect on follicle density and health (13) or a negative effect on both (14).

In this context, the present study set out to examine the effect of delayed processing and cryopreservation on ovarian tissue stored for fertility preservation using tissue processed in a clinically relevant manner. The sheep was used as a model organism for three reasons: 1) its structural similarity with human ovaries (15); 2) human ovarian tissue cryopreservation has been largely optimised using the sheep as a model; and 3) it is the only species, aside from rodents and primates, in which a live birth has been achieved after transplantation of frozen-thawed ovarian tissue (16). As the clinical value of cryopreserved ovarian tissue is largely determined by its potential to restore fertility following autologous grafting, xenotransplantation into immunodeficient mice was employed to examine the effect of delayed processing on *in vivo* development after cryopreservation and thawing. The aim of the study was to establish if processing delay of 24 or 48 h had an effect on primordial follicle morphology in fresh, frozen-thawed and xenotransplanted ovine ovarian cortical tissue.

## Materials and methods

### Tissue collection

Pairs of ovaries from six female lambs (*Ovis aries*, breed unknown) were collected from a local abattoir and transported on ice in 100 mL of transport medium [Leibovitz’s L-15 medium (Thermo Fisher, UK) supplemented with 100 U/mL penicillin and 100 μg/mL streptomycin (Sigma, UK) and 2.5 μg/mL amphotericin B (Sigma)]. Upon arrival at the laboratory, one ovary from each pair was immediately processed (no delay), while the other ovary was stored at 4°C in transport medium for 24 h or 48 h.

### Ovarian tissue processing and cryopreservation

Ovaries were processed for cryopreservation using procedures aligned with the relevant standard operating procedures (SOPs) applied at the Oxford Cell and Tissue Biobank (Oxford, UK), with some modification. Each ovary was bivalved and the ovarian medulla was gently dissected away using curved scissors under aseptic conditions. The outermost 1 mm of the ovarian cortex was cut to strips of approximately 5 × 2 × 1 mm using a surgical scalpel blade. All tissue handling was carried out in transport media on a *Medicool* ice block. Cortical strips were either fixed directly after processing (fresh samples) or cryopreserved. For cryopreservation, individual cortical strips were transferred to Nunc cryotubes and equilibrated in 1 mL of cryoprotectant medium containing L-15 supplemented with 1.5 M ethylene glycol (Sigma), 0.1 M sucrose (Sigma) and 3 mg/mL bovine serum albumin (BSA, Fisher Scientific, UK) for 1 h on ice. Cryopreservation was carried out in a controlled-rate freezer (IceCube 14S, SY-LAB, Sweden) using the following cooling programme: start temperature: 4°C; cooling rate I: −2°C/min to −9°C; cooling rate II: −0.3°C/min to −40°C; cooling rate III: −10°C/min to −140°C. Manual seeding was performed at −9°C and after freezing samples were stored in vapour phase liquid nitrogen.

### Tissue thawing

Cryotubes were thawed in a water bath at 30°C for 3 min, after which cortical strips were washed through three thawing solutions, containing a decreasing gradient of ethylene glycol (1 M, 0.5 M and 0 M, 0.1 M sucrose and 3 mg/mL BSA in L-15 medium, for 5 min each at room temperature. For samples that were to be xenotransplanted, 2.5 μg/ml amphotericin B, 100 U/ml penicillin and 100 μ g/ml streptomycin were added to the thawing solutions. Tissue was transplanted within 1 h or thawing.

### Mice

Immunodeficient female mice (SCID; CB17/Icr-Prkdc^SCID^/IcrIcoCrl) were obtained from Charles River Laboratories (Kent, UK) and housed together in a cage with filtered air supply under a 12:12 h light-dark cycle with *ad libitum* access to sterile food and water. The animals were acclimatised for two weeks prior to surgery and xenotransplantation was carried out at 8 weeks of age.

### Ethics approval

Animal work was carried out in accordance with project license number 30/3352 granted to Dr Suzannah Williams under The Animals (Scientific Procedures) Act 1986.

### Xenotransplantation

Thawed cortical strips (1 × 1 × 3 mm) were transplanted subcutaneously to the left flank of immunocompromised mice. Four strips were transplanted to each mouse with strips from one ovarian pair (no delay and 24 h delay) in two separate pockets created from the first incision and the second ovarian pair (no delay and 48 h delay) into two separate pockets created from the second incision. Each graft’s position was secured before the incision was closed. Mice were housed individually for the first 24 h after surgery. Recovery was monitored by observation and measuring body weight post-transplant; mice were weighed twice daily in the first 48 h and once daily thereafter. After a period of 10 days, the animals were sacrificed by cervical dislocation and all grafts retrieved.

### Histological analysis

Cortical strips were fixed in Bouin’s solution overnight at room temperature and embedded in paraffin wax. Serial sections were obtained at 5 µm thickness and slides were stained with haematoxylin and eosin (H&E). Samples were imaged at x400 magnification using a Leica DM2500 microscope. Images were captured using QCapture Pro 7 software. Follicles were graded in a blinded manner by two independent researchers, with a concordance rate of over 90%.

Primordial follicles were identified based on morphology defined by a single layer of flattened granulosa cells (17). Oocyte health was assessed as described previously (18), with slight modification. Oocytes were graded as healthy (grade 0) if they had no morphological markers, affected (grade 1) if they had either dense chromatin, eosinophilia or shrinkage of the ooplasm (one point each), degenerating (grade 2) if they had two of the above or nuclear pyknosis (two points) and atretic (grade ≥3) if they had more than two of the above markers (Figure 1). Up to 50 primordial follicles were analysed within each tissue fragment, except for xenotransplants, where all primordial follicles were included in the analysis.

**Figure 1.**
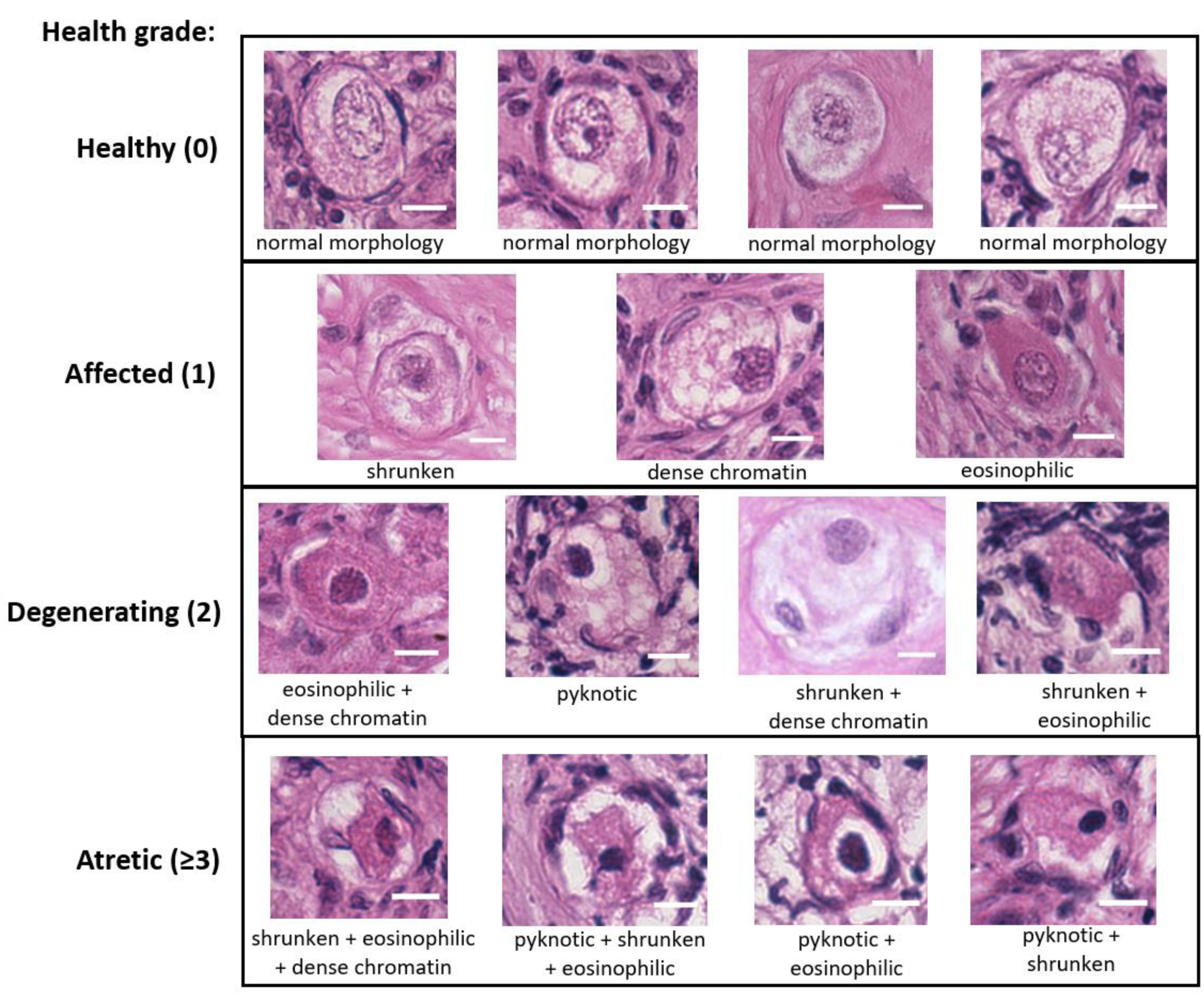
Health grading of primordial follicles based on morphological appearance. Bouin’s-fixed ovine ovarian tissue was analysed for the presence of primordial follicles (oocyte surrounded by a single layer of flattened pre-granulosa cells) using haematoxylin and eosin (H&E) staining. Follicles were scored based on the presence of morphological markers (dense chromatin, eosinophilia, shrunken ooplasm (one point each) or pyknosis (two points)) as healthy (0), affected (1), degenerating (2) or atretic (≥3). Scale bar: 10 μm.

### Data analysis

All statistical analyses were performed using R statistical software, version 3.5.0. A proportional odds model (clmm2; Christensen, 2015) was used to determine whether delay before processing affected follicle health, adjusting for individual sheep (random effect). A logistic regression model (glmer; Bates *et al*., 2015) was used to determine the odds of follicles being classified as growing after xenotransplantation, again adjusting for individual sheep. The total number of follicles per group (n) is indicated for each analysis. Statistical significance was defined as *p*<0.05. Data are presented as mean ± SEM or as odds ratios (ORs) with 95% confidence intervals (CI).

## Results

### The effect of processing delay on fresh ovarian tissue

Pairs of sheep ovaries were collected and either processed immediately (n=6) or stored at 4°C for 24 h (n=3) or 48 h (n=3) before processing. Primordial follicles were graded based on oocyte morphology as healthy (grade 0) or affected, degenerating or atretic (grades 1-3, hereafter referred to as ‘unhealthy’) based on the presence of nuclear condensation or pyknosis and shrinkage or eosinophilia of the ooplasm (Figure 1). The vast majority of follicles in all fresh tissues were healthy; 89.0±3.5% for the immediately processed (no delay) group, 90.6±1.7% for the 24 h-delay group and 79.2±8.2% for the 48 h-delay group (Figure 2A-C). Compared to immediately processed samples, a 24 h delay did not result in an increased likelihood of primordial follicles being graded as unhealthy (OR=0.78, 95%CI 0.38-1.60, *p>*0.05). In contrast, primordial follicles from samples processed 48 h after procurement were 2.5 times more likely to be graded as unhealthy compared to no delay samples (OR=2.47, 95%CI 1.29-4.71, *p*<0.01; Figure 2D).

**Figure 2.**
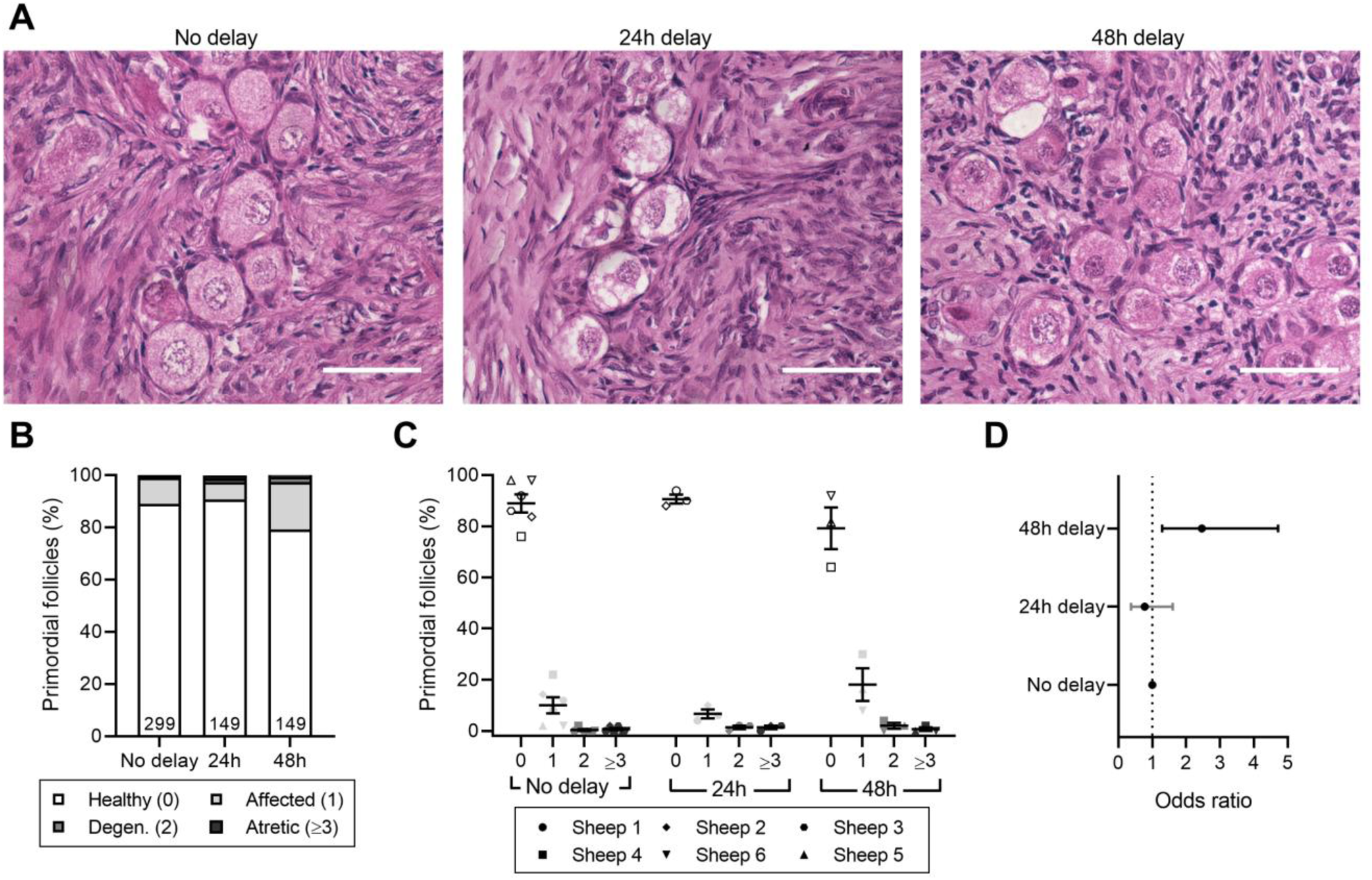
Effect of delayed processing on primordial follicle morphology in fresh ovarian tissue. Ovine ovarian tissue was either processed immediately (no delay; n=6), 24h (n=3) or 48h (n=3) after collection and fixed in Bouin’s solution. **A)** Representative images of fresh ovine ovarian tissue. Scale bar: 50 μm. **B)** Health distribution of primordial follicles in fresh ovine ovarian tissue. Numbers at the base of columns refer to the total number of follicles analysed. Primordial follicles (oocyte surrounded by a single layer of flattened pre-granulosa cells) were analysed using H&E staining and scored based on the presence of morphological markers (dense chromatin, eosinophilia, shrunken ooplasm (one point each) or pyknosis (two points)) as healthy (0), affected (1), degenerating (2) or atretic (≥3). **C)** Same data as in B, showing the distribution of health grades between animals (mean ± SEM). **D)** Proportional ORs and 95% CIs of follicles being graded as ≥ 1 when processed after 24h or 48h compared to the no delay group.

### The effect of processing delay on cryopreserved and thawed ovarian tissue

Consistent with current literature, cryopreservation caused damage to primordial follicles, with follicles in cryopreserved-thawed samples being over 28 times more likely to be graded as unhealthy compared to fresh tissue (OR=28.60, 95% CI 17.91-45.65, *p*<0.0001, Figure 3). Thus, the proportion of healthy primordial follicles was lower in cryopreserved-thawed tissue compared to fresh tissue, irrespective of processing time (Figure 4A-C). The mean proportion of healthy follicles in cryopreserved-thawed tissue with no processing delay was 26.6±17.0% and 20.4±2.1% for tissue processed 24 h after collection. Interestingly, tissue processed 48 h after collection had a higher proportion of healthy primordial follicles, 45.7±31.3%, though with considerable deviation between individuals (Figure 4C). This translated to a 44% decrease in the odds of a follicle being graded as unhealthy compared to immediately processed tissue (OR=0.56, 95% CI 0.36-0.87, *p*<0.01; Figure 4D). There was no difference in the health of primordial follicles after a 24 h delay before processing compared to immediately processed tissue (OR=0.93, 95% CI 0.61-1.41, *p*>0.05).

**Figure 3.**
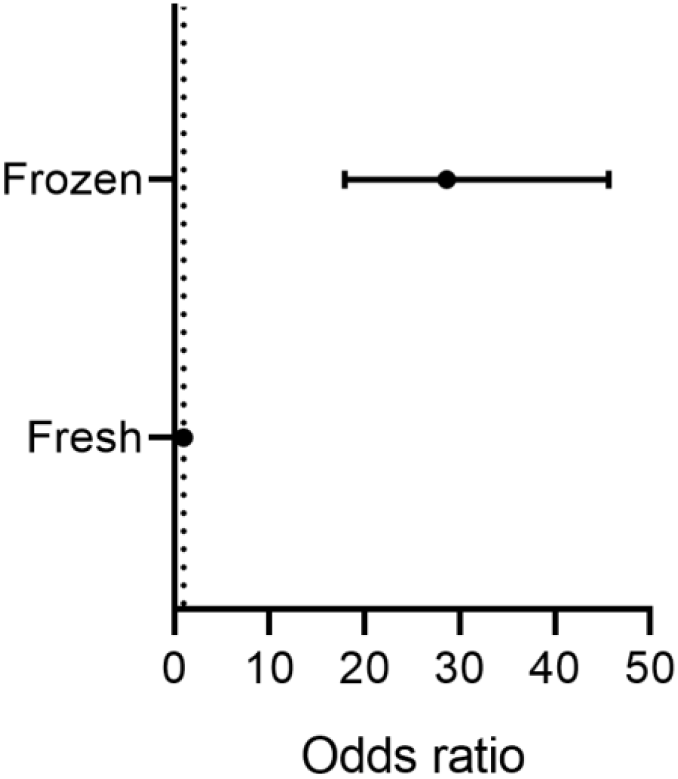
Effect of cryopreservation of ovarian tissue on primordial follicle morphology. Proportional ORs and 95% CIs of follicles being graded as ≥ 1 in fresh tissue compared to cryopreserved-thawed tissue. Ovine ovarian tissue was processed immediately and either fixed or cryopreserved-thawed and fixed (n=6) in Bouin’s solution. Primordial follicles (oocyte surrounded by a single layer of flattened pre-granulosa cells) were analysed using H&E staining and scored based on the presence of molecular markers (dense chromatin, eosinophilia, shrunken ooplasm (one point each) or pyknosis (two points)) as healthy (0), affected (1), degenerating (2) or atretic (3).Black CI bars indicate *p*<0.05.

**Figure 4.**
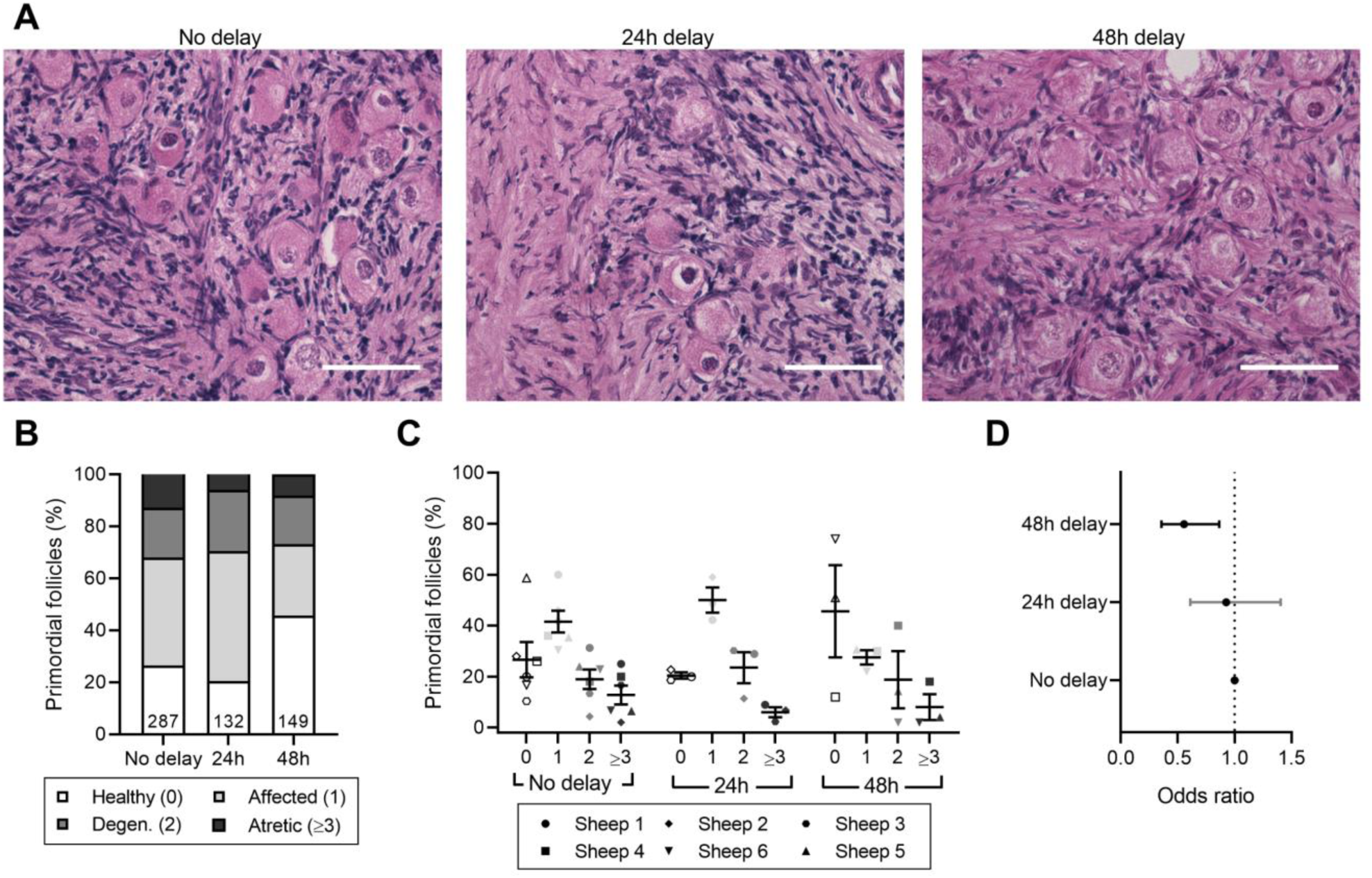
Effect of delayed processing on primordial follicle morphology in cryopreserved-thawed ovarian tissue. Ovine ovarian tissue was either processed immediately (no delay; n=6), 24h (n=3) or 48h (n=3) after collection, cryopreserved by slow freezing, thawed and fixed in Bouin’s solution. **A)** Representative images of cryopreserved-thawed ovine ovarian tissue. Scale bar: 50 μm. **B)** Health distribution of primordial follicles in cryopreserved-thawed ovine ovarian tissue. Numbers at the base of columns refer to the total number of follicles analysed. Primordial follicles (oocyte surrounded by a single layer of flattened pre-granulosa cells) were analysed using H&E staining and scored based on the presence of morphological markers (dense chromatin, eosinophilia, shrunken ooplasm (one point each) or pyknosis (two points)) as healthy (0), affected (1), degenerating (2) or atretic (≥3). **C)** Same data as in B), showing the distribution of health grades between animals (mean ± SEM). **D)** Proportional ORs and 95% CIs of follicles being graded as ≥ 1 when processed after 24h or 48h compared to the no delay group. Black CI bars indicate *p*<0.05.

### The effect of processing delay on ovarian tissue xenotransplanted after cryopreservation

Cryopreserved-thawed ovine ovarian tissue from 0 h, 24 h, and 48 h groups was xenografted subcutaneously into SCID mice for 10 days. The number of follicles found in each graft differed substantially across conditions; out of the six control grafts (immediately processed samples) only three contained follicles, with the remaining grafts containing potential degenerating follicle remnants (follicle-like structures with no oocyte; Table 1). Conversely, follicles were observed in two out of three grafts for the 24h-delay group and all three grafts for the 48h-delay group. Many primordial follicles were healthy after 10-day xenografting, however considerable variation was observed between grafts, particularly for the 48h-delay group. The proportion of healthy primordial follicles was 55.7±9.8% for the immediately processed group, 81.5±3.5% for the 24h-delay group and 43±28.8% for the 48h-delay group (Figure 5A-C). There was no difference in the likelihood of a primordial follicle being graded as unhealthy for the 24 h (OR=0.36, 95% CI 0.081-1.59, *p*>0.05) or 48 h delay groups (OR=5.0, 95% CI 0.80-30.83, *p*>0.05) compared to the no delay group (Figure 5D).

**Table 1.** Number of follicles per graft following 10-day xenotransplantation of cryopreserved-thawed ovine ovarian tissue with delayed processing.

| Animal | No delay | 24h delay | 48h delay |
| --- | --- | --- | --- |
| Sheep 1 | 25 follicles<br>(14 primordial, 56%) | 27 follicles<br>(14 primordial, 52%) | N/A |
| Sheep 2 | No follicles | 154 follicles<br>(86 primordial, 56%) | N/A |
| Sheep 3 | 6 follicles<br>(6 primordial, 100%) | No follicles | N/A |
| Sheep 4 | No follicles | N/A | 75 follicles<br>(50 primordial, 67%) |
| Sheep 5 | No follicles | N/A | 32 follicles<br>(16 primordial, 50%) |
| Sheep 6 | 2 follicles<br>(2 primordial, 100%) | N/A | 55 follicles<br>(26 primordial, 47%) |
| **Total** | **33 (22 primordial)** | **181 (100 primordial)** | **162 (92 primordial)** |

**Figure 5.**
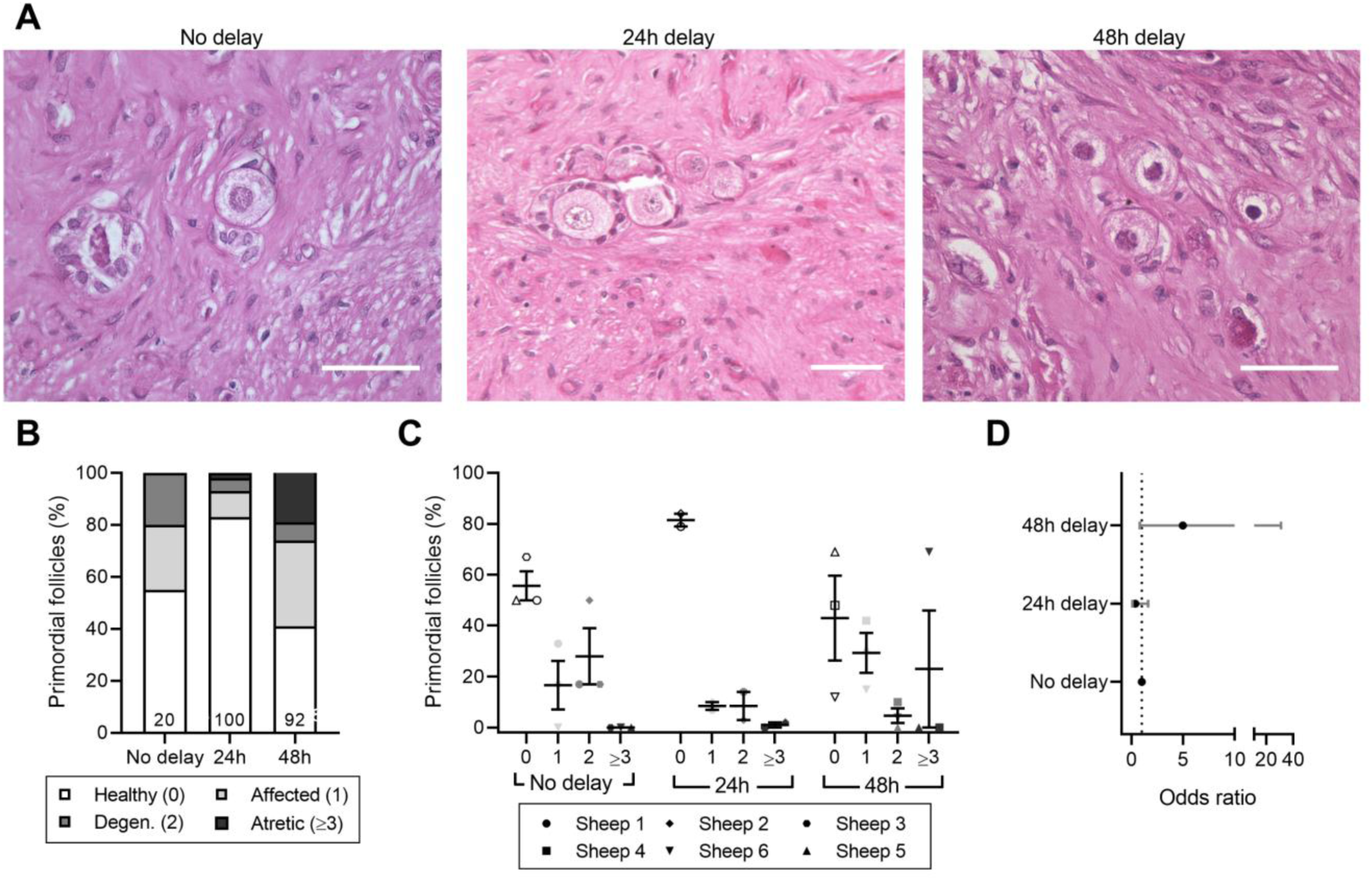
Effect of delayed processing on primordial follicle morphology in cryopreserved-thawed xenotransplanted ovarian tissue. Ovine ovarian tissue was either processed immediately (no delay; n=6), 24h (n=3) or 48h (n=3) after collection and cryopreserved. Samples were thawed and xenotransplanted subcutaneously for 10 days, after which they were fixed in Bouin’s solution. **A)** Representative images of cryopreserved-thawed ovine ovarian tissue. Scale bar: 50 μm. **B)** Health distribution of primordial follicles in cryopreserved-thawed ovine ovarian tissue. Numbers at the base of columns refer to the total number of follicles analysed. Primordial follicles (oocyte surrounded by a single layer of flattened pre-granulosa cells) were analysed using H&E staining and scored based on the presence of morphological markers (dense chromatin, eosinophilia, shrunken ooplasm (one point each) or pyknosis (two points)) as healthy (0), affected (1), degenerating (2) or atretic (≥3). **C)** Same data as in B), showing the distribution of health grades between animals (mean ± SEM). **D)** Proportional ORs and 95% CIs of follicles being graded as ≥ 1 when processed after 24h or 48h compared to the no delay group.

Of the eight grafts containing follicles, six contained both primordial and growing follicles indicating that follicle activation occurred during the xenografting period (Table 1; Figure 6). The other two grafts that contained follicles contained only primordial follicles and in very low numbers; 2 and 6 follicles (Table 1). The proportion of growing follicles was 17.3±30.0% for tissue processed immediately after collection, 46.0±2.8% for tissue processed after 24 h and 45.3±10.8% for tissue processed after 48 h (Figure 6A), with the highest variability between animals in the no delay group (Figure 6B). There was no difference in the likelihood of a follicle being classified as ‘growing’ between the groups (24 h delay OR=1.30, 95% CI 0.59-3.22, *p*>0.05; 48 h delay OR 1.26, 95% CI 0.55-3.96, *p*>0.05; Figure 6C).

**Figure 6.**
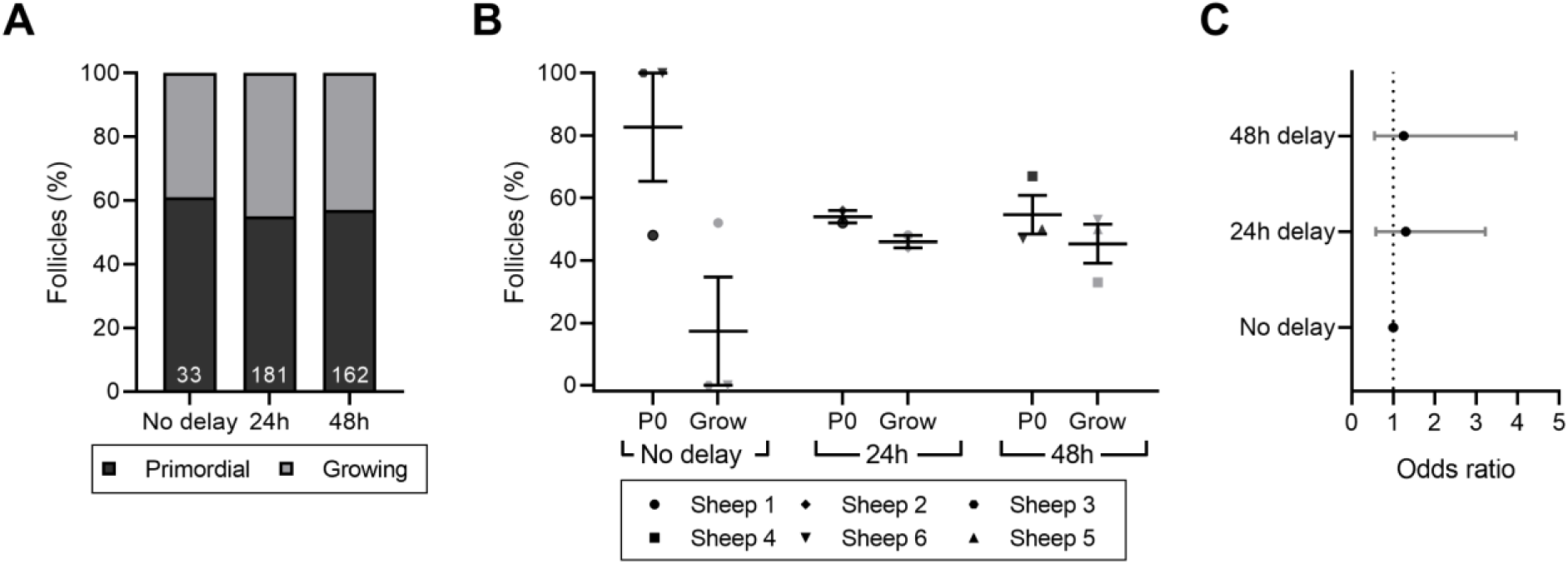
Follicle development after xenotransplantation of ovarian tissue with or without delayed processing. Ovine ovarian tissue was either processed immediately (no delay; n=6), 24h (n=3) or 48h (n=3) after collection and cryopreserved. Samples were thawed and xenotransplanted subcutaneously for 10 days, after which they were fixed in Bouin’s solution. **A)** Proportion of primordial and growing follicles in cryopreserved-thawed ovine ovarian tissue following 10-day xenotransplantation. Follicles were classified as primordial (P0; oocyte surrounded by a single layer of flattened pre-granulosa cells) or growing (Grow; oocyte surrounded by one or more layers of either a mixture of flattened and cuboidal granulosa cells, or a complete layer of cuboidal cells. **B)** Same data as in A), showing the distribution of health grades between animals (mean ± SEM). **C)** Proportional ORs and 95% CIs of follicles being classified as growing when processed after 24h or 48h compared to the no delay group.

## Discussion

This study demonstrates that a processing delay of 24 h at 4°C did not impact the health of the primordial follicles in fresh or cryopreserved ovine ovarian tissue. In contrast, a processing delay of 48 h resulted in primordial follicles being more likely to be graded as unhealthy compared to immediately-processed tissue, while tissue cryopreserved after 48 h was more likely to contain healthy follicles compared to the immediately-processed control. Xenografting of cryopreserved ovarian tissue confirmed tissue function in all experimental groups, with the delay in processing resulting in no difference in the health of primordial follicles, nor in the proportion of growing follicles.

Here we reported that cryopreservation had a significant negative impact on primordial follicle health, with only 20-40% of follicles being classified as healthy. Health classification criteria vary considerable between studies, and the criteria used in the present study were deliberately strict, in order to detect any discrete morphological changes within the follicles. Others have reported 50-80% healthy follicles following cryopreservation, which is consistent with our report if the classification criteria to judge the health of the follicles in this study is adjusted to match (21,22). We acknowledge that the results of the present study are somewhat limited by the use of morphology alone to assess primordial follicle health. However the criteria used were well-defined and in studies using a combination of morphological and molecular analyses, the two methods are generally in agreement (23). While the use of molecular markers may have complemented the results presented here, they present their own limitations including challenges in selecting an appropriate marker for the damage mechanism in question, in addition to the transience of some marker expression (24).

A 24 h processing delay had no significant effect on primordial follicle health post-cryopreservation and thawing. This finding is consistent with the presence of morphologically normal primordial follicles after a 20 h processing delay at 4°C of human ovarian cortical tissue as reported by Rosendahl *et al*. (12). Isachenko *et al*. also provided evidence that a 24 h cryopreservation delay at 4°C does not affect the proportion of morphologically normal follicles or follicle density in frozen-thawed ovarian cortical tissue (13). Conversely, another study described a decrease in the proportion of morphologically normal follicles induced by a 24 h freeze-delay prior to thawing and *in vitro* culture of human ovarian tissue (14), however, this study used vitrification, which relies on very high concentrations of cryoprotectants, while the present study employed slow freezing which is the currently accepted gold-standard method of ovarian tissue cryopreservation (25).

Remarkably, a processing delay of 48 h resulted in a 44% decrease in the probability of follicles being graded as affected, degenerating or atretic after cryopreservation, despite showing the opposite effect in fresh tissue. This is consistent with findings from Isachenko *et al*. who explored a 24h delay at 4°C suggesting a role for pre-cooling of human ovarian tissue before cryopreservation (13). It is possible that acclimatisation of the ovarian tissue at 4°C for 48 h provides a more gradual temperature transition, which primes the tissue to better respond to the challenges of cryopreservation and, thus, experience a reduced insult on the health of the primordial follicle pool. However, as there was significant variability in follicle health distribution between the three biological replicates in response to a 48 h delay, confirming this result with human ovarian tissue is required before such delays can be used for fertility preservation.

In this study we used xenotransplantation to determine if the delay in ovarian tissue cryopreservation had modified function. Importantly, all treatment groups contained grafts that supported follicle development. Nevertheless, there was significant variation in follicle numbers between grafts, with some grafts containing no follicles at all. The variable number of follicles in the graft was accounted for in the statistical model used. Total absence of follicles was observed in 3 out of 6 grafts that were processed immediately and 1 out of 3 24 h delay grafts, but all contained degenerating follicle remnants. Most surprisingly, delayed processing of up to 48 h did not adversely affect follicle development in cryopreserved-transplanted tissue, as primordial and growing follicles were observed in all three of the 48 h delay grafts. Thus the presence of follicles in 5 out of 6 grafts confirms that such a delay does not render the tissue deficient in follicles capable of growth nor does it hinder revascularisation of the tissue, a key component to the success of transplantation. By extension, this data also suggests that a processing delay of 24 or 48 h did not cause loss of function of the vasculature and the supporting stromal cells. This finding is consistent with insight from other studies that also identified follicles in ovarian tissue xenotransplanted after a 24 h delay in cryopreservation (12,13). Moreover, these results demonstrate that a processing delay of either 24 or 48 h had no significant impact on primordial follicle health nor on the proportion of growing follicles after xenotransplantation indicating that delayed processing does not affect the potential for follicle development following transplantation. To the best knowledge of the authors, the work presented here is the first report of successful ovarian tissue grafting after a 48 h delay in cryopreservation.

Although cryopreservation of ovarian tissue with as little delay as possible seems obvious, and is considered the gold standard in most countries (29), it is possible that transient cold ischaemia during delayed processing induces a metabolic adjustment which enables quiescent primordial follicles to better withstand ischaemia prior to revascularisation. Energetic adjustment to ischaemia is well-documented in muscle tissue (30) and it is possible that the oocyte is also equipped with the capacity to adjust to metabolism in avascular conditions. Furthermore, there is some evidence to suggest that ischaemic preconditioning can improve the outcomes of organ transplantation (31) and in this context, our findings might suggest that the benefits of ischaemic preconditioning are not lost after cryopreservation and thawing of the tissue. This, taken together with studies using human tissue, which demonstrated that follicles can survive a processing delay of delay of up to 20 h at 4°C and lead to a live birth (8,12,13), indicates that ovarian tissue responds to delayed cryopreservation favourably without a negative impact on the primordial pool.

The results we present here have provided strong evidence to support the idea that ovarian tissue can be chilled for 24 or even 48 h prior to cryopreservation with no net negative impact on the health of the primordial pool of follicles after transplantation and *in vivo* development of the tissue. This lends strength to an emerging body of evidence suggesting the delayed processing is safe, or even beneficial, for cryopreserved ovarian tissue. Validation of these findings using human ovarian tissue or other endangered species of interest would thereby present a new, feasible approach to increasing access to ovarian tissue freezing for fertility preservation. It should be noted that in the present study ovaries were left intact until the time of processing, therefore, it cannot be concluded that ovarian biopsies would have a similar response. Nevertheless, these findings are of high relevance to fertility preservation and, in particular, the way ovarian tissue cryopreservation is organised on a local and national level for humans and endangered species. An update of the current guidelines to reflect an extended window for tissue processing will ensure that the modality is more widely available to patients without the need to create new facilities. Fundamentally, such an advancement is a great step towards ensuring that every cancer patient who has a need, has an option.

## Acknowledgements

The Authors wish to thank all staff of the Oxford Cell and Tissue Biobank (OCTB) for their assistance with training in tissue processing and cryopreservation, Prof. David Bennett for use of the surgery facilities, and all staff of Biomedical Sciences John Radcliffe for assistance with the mice used in this study.

## References

1. Miller KD, Siegel RL, Lin CC, Mariotto AB, Kramer JL, Rowland JH, et al. Cancer Treatment and Survivorship Statistics, 2016. 2016;66(4):271–89.

2. Phillips SM, Padgett LS, Leisenring WM, Stratton KK, Bishop K, Krull KR, et al. Survivors of childhood cancer in the United States: Prevalence and burden of morbidity. Cancer Epidemiol Biomarkers Prev. 2015;24(4):653–63.

3. Armstrong GT, Chen Y, Yasui Y, Leisenring W, Gibson TM, Mertens AC, et al. Reduction in late mortality among 5-year survivors of childhood cancer. N Engl J Med. 2016;374(9):833–42.

4. Spoudeas HA. Growth and endocrine function after chemotherapy and radiotherapy in childhood. Eur J Cancer. 2002;38(13):1748–59.

5. Duffy C, Allen S. Medical and psychosocial aspects of fertility after cancer. Cancer J. 2009;15(1):27–33.

6. Donnez J, Dolmans M-M. Fertility Preservation in Women. N Engl J Med. 2017;377(17):1657–65.

7. Gellert SE, Pors SE, Kristensen SG, Bay-Bjørn AM, Ernst E, Yding Andersen C. Transplantation of frozen-thawed ovarian tissue: an update on worldwide activity published in peer-reviewed papers and on the Danish cohort. J Assist Reprod Genet. 2018;

8. Dittrich R, Lotz L, Keck G, Hoffmann I, Mueller A, Beckmann MW, et al. Live birth after ovarian tissue autotransplantation following overnight transportation before cryopreservation. Fertil Steril. 2012;97(2):387–90.

9. Jensen AK, Macklon KT, Fedder J, Ernst E, Humaidan P, Andersen CY. 86 successful births and 9 ongoing pregnancies worldwide in women transplanted with frozen-thawed ovarian tissue: focus on birth and perinatal outcome in 40 of these children. J Assist Reprod Genet. 2017;34(3):337.

10. Majno G, Joris I. Apoptosis, oncosis, and necrosis: An overview of cell death. Am J Pathol. 1995;146(1):3–15.

11. Kim SS, Battaglia DE, Soules MR. The future of human ovarian cryopreservation and transplantation: Fertility and beyond. Fertil Steril. 2001;75(6):1049–56.

12. Rosendahl M, Schmidt KT, Ernst E, Rasmussen PE, Loft A, Byskov AG, et al. Cryopreservation of ovarian tissue for a decade in Denmark: A view of the technique. Reprod Biomed Online. 2011;22(2):162–71.

13. Isachenko V, Todorov P, Isachenko E, Rahimi G, Tchorbanov A, Mihaylova N, et al. Long-time cooling before cryopreservation decreased translocation of phosphatidylserine (Ptd-L-Ser) in human ovarian tissue. PLoS One. 2015;10(6):1–14.

14. Klocke S, Tappehorn C, Griesinger G. Effects of supra-zero storage on human ovarian cortex prior to vitrification-warming. Reprod Biomed Online. 2014;29(2):251–8.

15. Oktay K. Ovarian tissue cryopreservation and transplantation: Preliminary findings and implications for cancer patients. Hum Reprod Update. 2001;7(6):526–34.

16. Gosden RG, Baird DT, Wade JC, Webb R. Restoration of fertility to oophorectomized sheep by ovarian autografts stored at-196°c. Hum Reprod. 1994;9(4):597–603.

17. Lundy T, Smith P, O’Connell A, Hudson NL, McNatty KP. Populations of granulosa cells in small follicles of the sheep ovary. J Reprod Fertil. 1999;115(2):251–62.

18. Walker CA, Bjarkadottir BD, Fatum M, Lane S, Williams SA. Variation in follicle health and development in cultured cryopreserved ovarian cortical tissue: a study of ovarian tissue from patients undergoing fertility preservation. Hum Fertil. 2019 May 23;0(0):1–11.

19. Christensen RHB. ordinal-Regression Models for Ordinal Data. R package version 2015.6-28. See http://www.cranr-project.org/package=ordinal. 2015;

20. Bates D, Mächler M, Bolker B, Walker S. Fitting Linear Mixed-Effects Models Using lme4. J Stat Softw. 2015;67(1):201–10.

21. Faustino LR, Santos RR, Silva CMG, Pinto LC, Celestino JJH, Campello CC, et al. Goat and sheep ovarian tissue cryopreservation: Effects on the morphology and development of primordial follicles and density of stromal cell. Anim Reprod Sci. 2010;122(1–2):90–7.

22. Cecconi S, Capacchietti G, Russo V, Berardinelli P, Mattioli M, Barboni B. In Vitro Growth of Preantral Follicles Isolated from Cryopreserved Ovine Ovarian Tissue. Biol Reprod. 2004;70(1):12–7.

23. Talevi R, Sudhakaran S, Barbato V, Merolla A, Braun S, Nardo M Di, et al. Is oxygen availability a limiting factor for in vitro folliculogenesis? PLoS One. 2018;13(2):e0192501.

24. Pampanini V, Wagner M, Asadi-Azarbaijani B, Oskam IC, Sheikhi M, Sjödin MOD, et al. Impact of first-line cancer treatment on the follicle quality in cryopreserved ovarian samples from girls and young women. Hum Reprod [Internet]. 2019 Sep 29;34(9):1674–85. Available from: https://academic.oup.com/humrep/article/34/9/1674/5549608

25. Dolmans MM, Manavella DD. Recent advances in fertility preservation. J Obstet Gynaecol Res. 2019;45(2):266–79.

26. Lane BR, Russo P, Uzzo RG, Hernandez A V., Boorjian SA, Thompson RH, et al.c Comparison of cold and warm ischemia during partial nephrectomy in 660 solitary kidneys reveals predominant role of nonmodifiable factors in determining ultimate renal function. J Urol. 2011;185(2):421–7.

27. Van Eyck AS, Bouzin C, Feron O, Romeu L, Van Langendonckt A, Donnez J, et al. Both host and graft vessels contribute to revascularization of xenografted human ovarian tissue in a murine model. Fertil Steril. 2010;93(5):1676–85.

28. Van Eyck AS, Jordan BF, Gallez B, Heilier JF, Van Langendonckt A, Donnez J. Electron paramagnetic resonance as a tool to evaluate human ovarian tissue reoxygenation after xenografting. Fertil Steril. 2009;92(1):374–81.

29. Donnez J, Dolmans MM, Pellicer A, Diaz-Garcia C, Sanchez Serrano M, Schmidt KT, et al. Restoration of ovarian activity and pregnancy after transplantation of cryopreserved ovarian tissue: A review of 60 cases of reimplantation. Fertil Steril. 2013;99(6):1503–13.

30. Lanza IR, Wigmore DM, Befroy DE, Kent-Braun JA. In vivo ATP production during free-flow and ischaemic muscle contractions in humans. J Physiol. 2006;577(1):353–67.

31. Ambros JT, Herrero-Fresneda I, Borau OG, Boira JMG. Ischemic preconditioning in solid organ transplantation: From experimental to clinics. Transpl Int. 2007;20(3):219–29.

